# SOX9 is required for kidney fibrosis and regulates NAV3 to control renal myofibroblast function in mice and humans

**DOI:** 10.1101/838441

**Authors:** Sayyid Raza, Elliot Jokl, James Pritchett, Katherine Martin, Kim Su, Kara Simpson, Aoibheann F Mullan, Varinder Athwal, Daniel T Doherty, Leo Zeef, Neil C Henderson, Philip A Kalra, Neil A Hanley, Karen Piper Hanley

## Abstract

Renal fibrosis is a common endpoint for many chronic kidney diseases. Extracellular matrix (ECM) from myofibroblasts causes progressive scarring and organ failure. The mechanisms underlying fibrogenesis and how it is sustained are incompletely understood. Here, we show that the transcription factor, Sex determining region Y-box 9 (SOX9), is required for kidney fibrosis. From genome-wide analysis we identify Neuron navigator 3 (NAV3) downstream of SOX9. NAV3 was upregulated in kidney disease in patients and following renal injury in mice colocalised with SOX9. By establishing an in vitro model of renal pericyte transition to myofibroblast we demonstrated that NAV3 is required for multiple aspects of fibrogenesis including actin polymerization linked to cell migration and sustaining SOX9 and active YAP1 levels. In summary, our work discovers novel SOX9-NAV3-YAP1/SOX9 circuitry as a new mechanism to explain the progression of kidney fibrosis and points to NAV3 as a novel target for pharmacological intervention.

## Introduction

Fibrotic scarring is common to many chronic kidney diseases (CKD). It is characterized by the pathological deposition of collagen-rich extracellular matrix (ECM) proteins from profibrotic (‘activated’) myofibroblasts in response to kidney injury and leads to loss of renal function ^1-3^. The fibrotic ECM increases organ stiffness. This rigidity perpetuates disease progression ^4, 5^. Despite renal fibrosis being a significant contributor to patient morbidity and mortality there are no anti-fibrotic drugs currently approved. In large part this reflects limited understanding of the underlying mechanisms, against which new therapies might be targeted, including how renal myofibroblasts arise. Fibrotic ECM signals via cell-surface integrins to control the cytoskeletal and mechanical properties of myofibroblasts ^4, 6^. Previously, we identified cytoskeletal alterations in liver myofibroblasts, downstream of integrin β1, mediated through actomyosin signaling via P21-activated kinase (PAK1) and the mechanical transcriptional effector, Yes-associated protein 1 (YAP1) ^7, 8^. In kidney, similar mechanisms have been invoked where YAP1 upregulates TGFβ, a potent profibrotic cytokine, to further activate myofibroblasts ^9, 10^. Pharmacological inhibition of YAP1 has partly ameliorated fibrosis in both kidney ^10, 11^ and liver ^7, 8^. These findings suggest value from considering mechanisms that might be shared across organs.

In liver, the transcription factor Sex determining region Y-box 9 (SOX9), downstream of TGFβ and YAP1, promotes fibrosis in vivo and in vitro by upregulating a wide array of genes encoding pathological ECM components ^8, 12-15^. SOX9 levels predict fibrosis disease progression within four years ^8^. The concentration of circulating proteins encoded by SOX9-target genes, such as osteopontin and vimentin, also correlates with the severity of liver fibrosis ^12^. SOX9 is known to be important in kidney development in mouse ^16, 17^and human ^18^. While it has been detected in renal fibrosis ^19^, SOX9’s role in vivo in renal fibrosis has been unaddressed and any putative mechanism of action unidentified. How myofibroblast activation is perpetuated in progressive renal fibrosis is unknown.

In this study, we hypothesized that if SOX9 was required for renal fibrosis in vivo, this would unlock an opportunity to discovery critical downstream target genes responsible for the profibrotic phenotype of renal myofibroblasts. We studied chronic kidney disease in patients and undertook SOX9 inactivation in the context of unilateral ureteric obstruction (UUO) in mouse to induce renal injury and fibrosis. We used transcriptomic deep sequencing (RNAseq) to pinpoint clusters of SOX9 target genes in vivo. We discovered and investigated Neuron navigator 3 (NAV3) as a target of SOX9 and novel mediator of myofibroblast function in kidney fibrosis. This included establishing a new model of pericyte activation into myofibroblasts. Together, the data prove the requirement for a critical transcription factor in promoting renal fibrosis, describe a new mechanism of action, and discover a novel target, NAV3, for therapeutic consideration.

## Results

### SOX9 is expressed in myofibroblasts in chronic kidney disease in human and mouse

By immunohistochemistry nuclear SOX9 was detected in chronic interstitial nephritis consistent with previous reports of the transcription factor in regenerative tubular cells ^20, 21^ (Figure 1). In diabetic and membranous nephropathies, SOX9 could also be observed in some nuclei within the glomerulus (Figure 1 and Supplementary Figure 1). In addition, the transcription factor was apparent in the nuclei of elongated myofibroblasts positive for α-Smooth muscle actin (α-SMA). These cells localized to areas of scarring in chronic interstitial nephritis and surrounding the glomerulus in Bowman’s capsule in diabetic nephropathy (Figure 1). SOX9 also localized to similar α-SMA-positive regions in focal segmental glomerulosclerosis and membranous nephropathy (Supplementary Figure 1). This pattern and colocalization with α-SMA is indicative of SOX9 in renal myofibroblasts in patients with chronic kidney disease.

**Figure 1.**
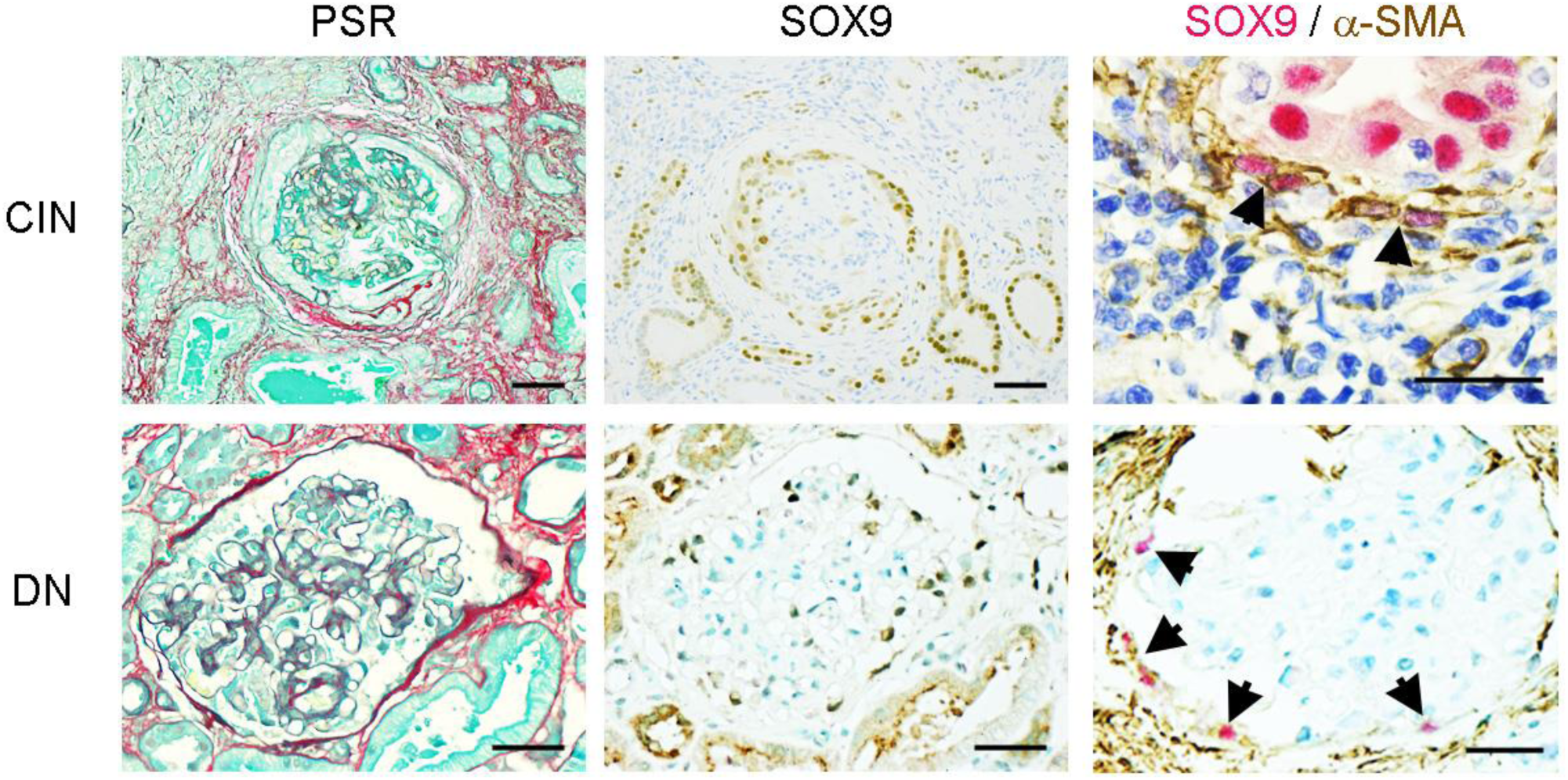
Localization of SOX9 in chronic kidney disease in patients. Serial kidney sections from chronic interstitial nephritis (CIN; top panel) and diabetic nephropathy (DN; bottom panel) show collagen deposition by picrosirius red (PSR, red, left) staining, or immunohistochemistry for SOX9 (brown, middle), and SOX9 (red) and α-SMA (brown) (right). Arrowheads point to myofibroblasts stained with cytoplasmic α-SMA and nuclear SOX9. Size bars = 25 μm.

UUO is the established model of renal injury in rodents which, after two weeks, causes profound ipsilateral interstitial fibrosis. The contralateral kidney is unaffected. Nuclear SOX9 was clearly visible at two weeks in the ipsilateral tubular cells post-UUO in mouse consistent with a regenerative response (Figure 2a-b). Virtually identical to patients with chronic kidney disease, SOX9 was also detected in spindle-shaped α-SMA (and PDGFRβ) positive cells in the interstitium surrounding tubules and glomeruli, including Bowman’s capsule (Figure 2c-f). Because some non-specific extracellular staining was observed in the brush border in the unaffected contralateral kidney, we corroborated our findings with *in situ* hybridization (ISH) for *Sox9* (Figure 2d). Collectively, these data confirm that the distribution of SOX9 in myofibroblasts during renal fibrosis is the same in human and mouse, validating inactivation studies in mouse to discover SOX9 function *in vivo*.

**Figure 2.**
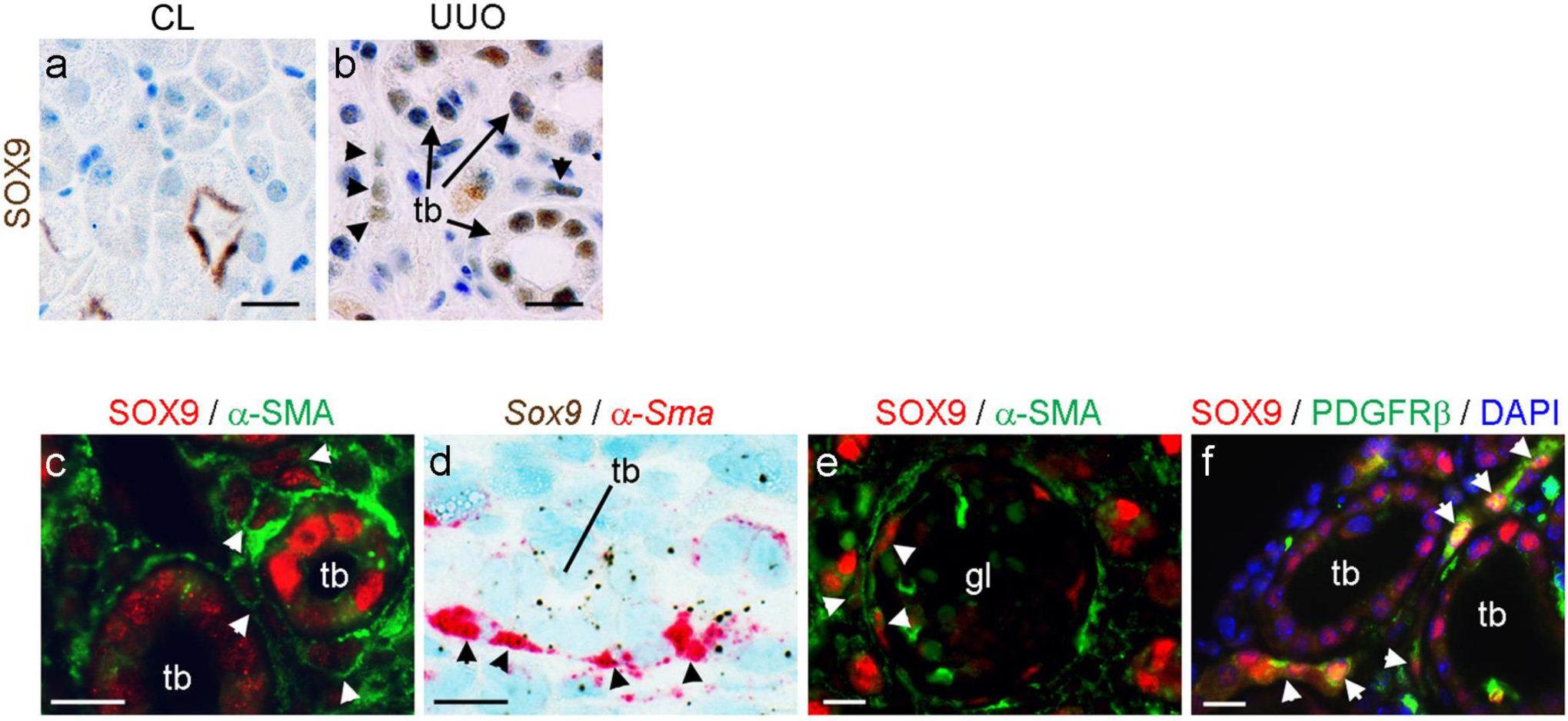
SOX9 becomes detected in ipsilateral (fibrotic) kidney following UUO in mice. (**a**) Non-specific staining in the brush border of kidney tubules in the unaffected contralateral (CL) kidney using the anti-SOX9 antibody. (**b**) Nuclear SOX9 detection in the ipsilateral kidney following UUO in tubules (tb; arrows) and interstitial cells (arrowheads). (**c-f**) Colocalization of SOX9 protein (c, e, and f) or transcript (d) with α-SMA (c-e) or PDGFRβ (counterstained with DAPI) in the ipsilateral kidney following UUO. Arrowheads point to colocalization in interstitial cells or cells in Bowman’s capsule. Tb, tubule; gl, glomerulus. Size bars = 10 μm.

### SOX9 loss-of-function improves kidney fibrosis

Widespread deletion of *Sox9* during development is lethal in mouse. To overcome this we used an inducible model to inactivate *Sox9* using Tamoxifen (Tam) in Sox9^fl/fl^;RosaCreER^+/−^ adult mice with Tam-treated Sox9^fl/fl^;RosaCreER^−/−^ mice as control (termed ‘*Sox9* null’ and ‘*Sox9* control’ respectively from hereon) ^8^. Tam was injected three times in the week prior to UUO. Fibrosis was induced in the ipsilateral kidney over 10 days and compared to the unaffected contralateral kidney. Tam led to complete loss of nuclear SOX9 in the kidney without compromising mouse survival (Supplementary Figure 2). Post-UUO, ipsilateral fibrosis was markedly improved by SOX9 loss. Collagen deposition by picrosirius red (PSR) staining and myofibroblast accumulation marked by α-SMA and PDGFRβ staining were approximately halved compared to renal fibrosis post-UUO in *Sox9* control tissue (Figure 3a-d). Similar data were apparent from whole kidneys at the level of transcription (Figure 3e-h). These data indicate that SOX9 is required for renal fibrosis in mice following ureteric ligation.

**Figure 3.**
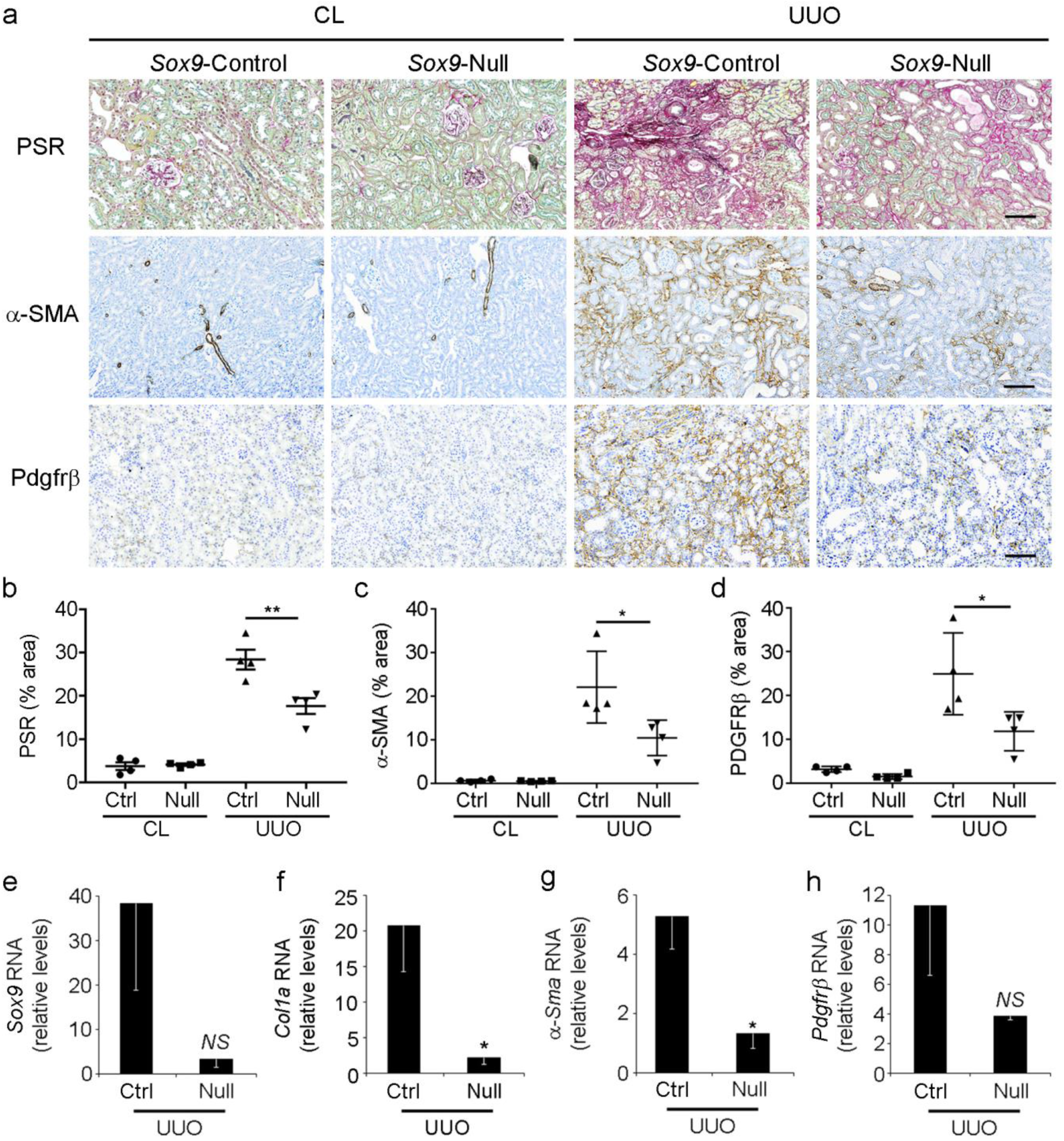
SOX9 loss reduces fibrosis and myofibroblast accumulation in the ipsilateral kidneys of mice following UUO. (**a**) Picrosirius red (PSR) staining for collagen deposition (top, red) and immunohistochemistry counterstained with toluidine blue for α-SMA (middle, brown) or PDGFRβ (bottom, brown) in the ipsilateral (UUO) and contralateral (CL) kidneys post-UUO in *Sox9* null and *Sox9* control mice. Size bars = 200 μm. (**b-d**) Quantification of surface area in UUO and CL kidneys post-UUO in *Sox9* control (Ctrl) and *Sox9* null (Null) mice covered by (b) PSR, (c) α-SMA or (d) PDGFRβ. (**e-h**) Relative RNA levels from whole UUO kidney in *Sox9* null or *Sox9* control mice. Data in the charts show means ± s.e.m. (n=4 mice for each group). *P<0.05, **P<0.01. NS, not significant.

### Delineating genome-wide pathways downstream of SOX9 in kidney fibrosis identifies *Nav3*

We wanted to discover what profibrotic genes were regulated by SOX9. Reassured that whole kidney analysis had detected clear differences for our user-defined genes, we undertook RNAseq in biological replicate following UUO on ipsilateral versus contralateral kidneys in *Sox9* control and *Sox9* null mice. Following hierarchical analysis, differentially expressed genes clustered into four categories on heatmap (Figure 4a, Supplementary Figure 3 and RNAseq dataset E-MTAB-8429). In the second largest category, Cluster 3, loss of SOX9 returned gene expression that had been upregulated by fibrosis back to control levels. Cluster 3 from Figure 4a is expanded in Supplementary Figure 4. By gene ontology, these 168 genes were heavily linked to fibrosis and ECM (Supplementary Figure 5). Within the most highly ranked disease and functional category (‘Cancer, connective tissue disorders, organismal injury and abnormalities’) we identified *Nav3*, which was one of the fifteen most differentially expressed genes in the entire dataset (log2 fold change, −2.98 in response to SOX9 inactivation) (Figure 4b and Supplementary Table 1). We verified the RNAseq findings by direct RT-qPCR. *Nav3* transcription was increased in the fibrotic kidney post-UUO but returned to baseline levels when Sox9 was absent (Figure 4c).

**Figure 4.**
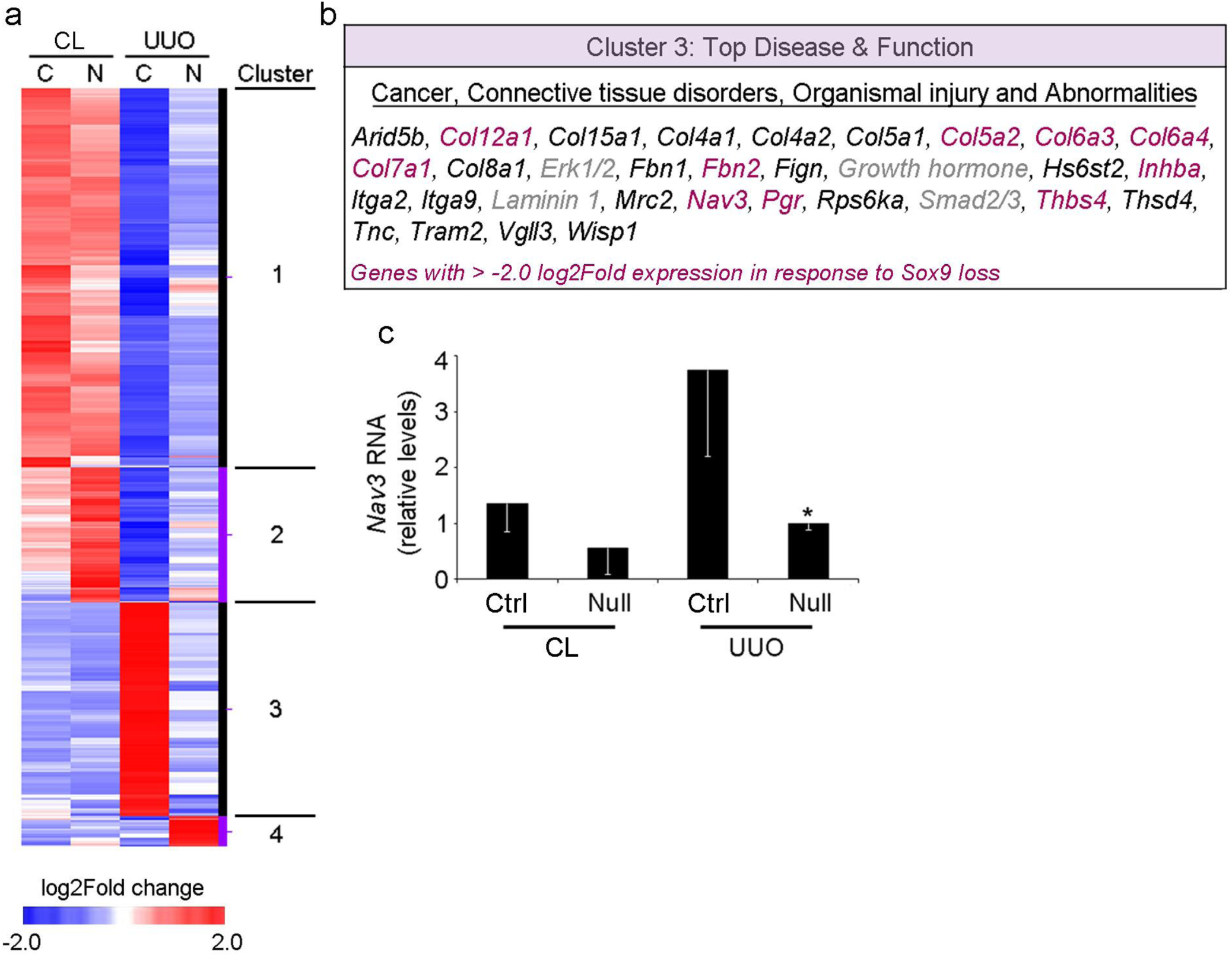
Profiling SOX9-dependent genes genome-wide in renal fibrosis identifies NAV3. (**a**) Cluster analysis and heatmap of mean gene expression changes (P<0.05) in whole contralateral (CL, unaffected) or ipsilateral (UUO) kidneys from *Sox9* null (N) and control (C) mice following UUO. Data for individual replicates are shown in Supplementary figure 3. Four clusters were identified based on upregulated (red) and downregulated (blue) gene expression. Individual genes from Cluster 3 (upregulated in renal fibrosis, SOX9-dependent) are shown in Supplementary figure 4. (**b**) Top disease function from gene ontology analysis of Cluster 3 (the small number of genes in grey are part of the category but unidentified in Cluster 3). (**c**) RTqPCR quantification for *Nav3* from whole contralateral (CL, unaffected) or ipsilateral (UUO) kidneys from *Sox9* null and control (Ctrl) mice following UUO. Mean ± s.e.m, n=4 mice for each group, *P<0.05.

### NAV3 is upregulated in kidney fibrosis

NAV3 has not been connected previously to any form of organ fibrosis or myofibroblast function. In normal mouse kidney, minimal NAV3 expression could be detected in and around vasculature adjacent to glomeruli (Figure 5a-c). In keeping with the distribution of SOX9 in fibrotic kidney post-UUO, NAV3 became detected in tubular cells and some cells of Bowman’s capsule; as well as spindle-shaped interstitial cells surrounding tubules and glomeruli (Figure 5a-d). We confirmed colocalization with *Sox9* and *α-Sma* by ISH (Figure 5d). NAV3 detection was the equivalent in kidney tissue from patients with multiple etiologies of CKD (Figure 5e-h). We quantified *Nav3* transcripts in fibrotic mouse kidney by ISH in the presence or absence of SOX9. Total *Nav3* was halved by loss of SOX9 and, in particular, appeared missing in the interstitial region in keeping with the reduced interstitial scarring and myofibroblast accumulation in *Sox9* null mice subjected to UUO (Figure 5i-j). We identified three conserved SOX9 binding sites (BS) surrounding the *NAV3* locus (Supplementary Figure 6a-b). By transient transfection and luciferase assay, inactivation of BS2 and 3 but not BS1 diminished reporter gene activity (Supplementary Figure 6c-e). Taken together, these data indicate that *NAV3* is dependent on and colocalizes with SOX9 in mouse and human renal fibrosis.

**Figure 5.**
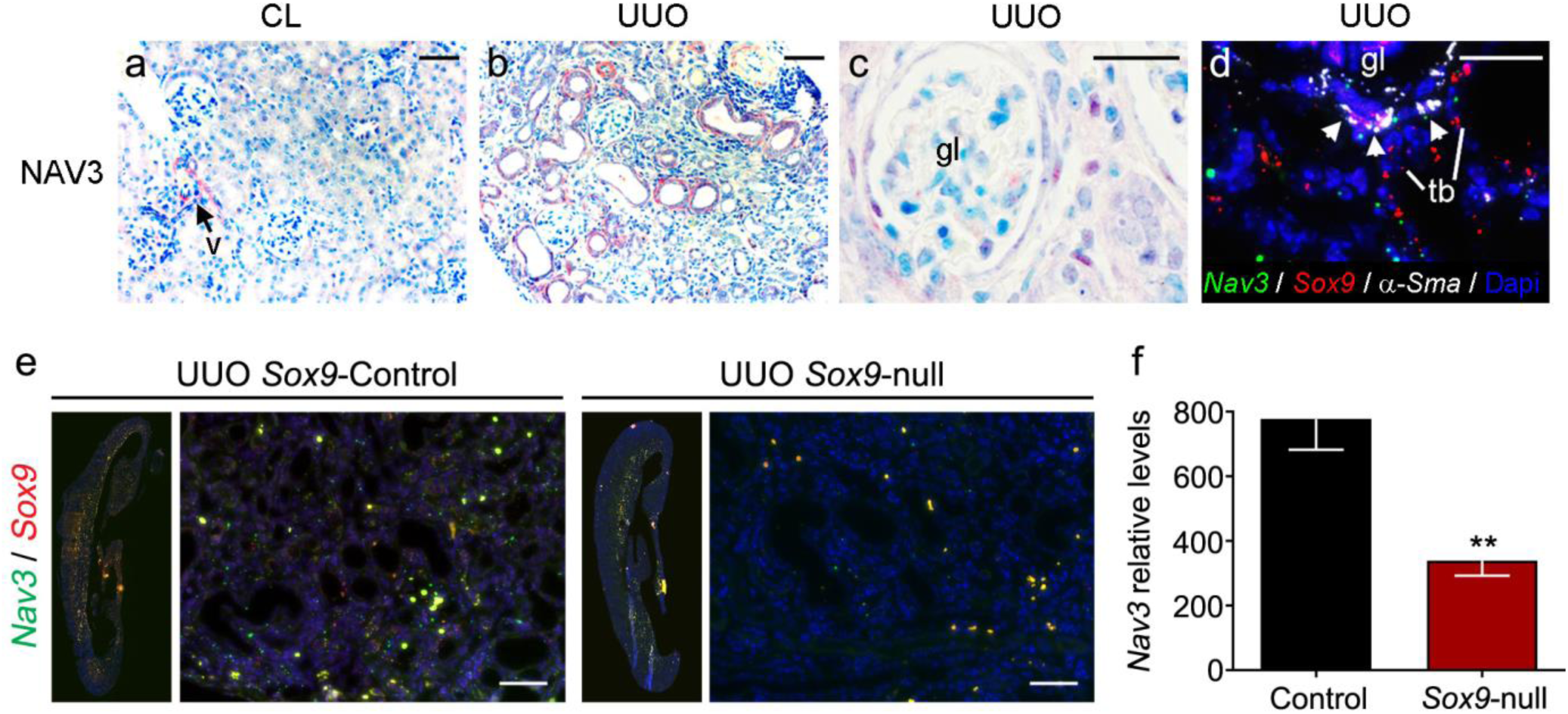
NAV3 localization in renal fibrosis in mice and human. Immunohistochemistry (red) counterstained with toluidine blue (**a-c**) or in situ hybridization (ISH; **d**) for NAV3 in mouse kidney. NAV3 localized around the vasculature (arrow) in (**a**) unaffected contralateral (CL) kidney and (**b-c**) ipsilateral fibrotic kidney following UUO (UUO; *Sox9* control mice). (**b**) NAV3 is detected in tubules post-injury (equivalent to SOX9 in Figure 2b). (**c**) NAV3 is also detected in the interstitial regions surrounding glomeruli (gl) and in Bowman’s capsule. (**d**) Multiplex in situ hybridisation colocalizes transcripts for *Nav3* (green), *Sox9* (red) and *α-Sma* (white) in interstitial cells (arrowheads) between the glomerulus and tubules (tb). (**e-h**) Immunohistochemistry for NAV3 (red) counterstained with toluidine blue in renal fibrosis due to chronic kidney disease in patients: chronic interstitial nephritis (CIN; **e-f**), diabetic nephropathy (DN; **g**) and hypertensive nephropathy (HN; **h**). (**e**-**f**) NAV3 localized to tubular cells in CIN as well as to Bowman’s capsule surrounding the glomerulus (gl; arrowheads). NAV3 was detected in interstitial regions between tubules (tb) in all cases (e.g. arrowheads in **h**). (**i-j**) Transcript localisation and quantification for *Sox9* (red) and *Nav3* (green) by ISH in *Sox9* control and *Sox9* null kidneys following UUO-induced fibrosis. Example images shown from the whole kidney and magnified area. (**j**) Quantification of *Nav3* transcripts by surface area in ipsilateral UUO kidneys from *Sox9* control and *Sox9* null mice. Mean ± s.e.m, **P<0.01. Size bars = 50 μm (a-d) and 25 μm (e-i).

### NAV3 is required by activated renal pericytes for cell migration and myofibroblast function

Myofibroblasts and the mechanosensitive transcription factor, YAP1, downstream of profibrotic TGFβ, are central to interstitial renal fibrosis ^10, 11, 22^. This is directly analogous to liver fibrosis, which in turn is dependent on the hepatic stellate cell (HSC). The HSC is a pericyte that when quiescent surrounds endothelial cells and upon activation migrates into areas of damage to lay down fibrotic ECM. HSC activation has been extensively studied in vitro by isolating quiescent cells by density gradient and culturing them on tissue culture plastic. We adapted this model to extract primary pericytes from normal mouse kidney. Directly comparable to their hepatic counterparts, on day 1 cells were rounded and lacked organized cytoplasmic α-SMA or nuclear SOX9 (Figure 6a). However, after culture for 14 days, again identical to activated HSCs, the pericytes had transitioned into a myofibroblast phenotype with filamentous α-SMA and robust nuclear detection of SOX9 (Figure 6a). We adopted the terms quiescent renal pericytes (QuiRPs) and activated renal pericytes (ARPs) for these two states. ARPs responded to TGFβ by increasing levels of SOX9, α-SMA and COL1 protein (Figure 6b). This new model allowed us to model renal fibrogenesis in vitro and study intracellular NAV3 function. NAV3 protein was barely detected in QuiRPs but strongly detected in ARPs when it organized linearly at the tips of actin filaments in the cell periphery, consistent with a role stabilizing microtubules in breast cancer cells (Figure 6c) ^23^. Abrogating SOX9 in ARPs by siRNA reduced levels of *α-Sma* and *Nav3*, further demonstrating the dependence of *Nav3* transcription on SOX9 (Figure 6d). We inactivated NAV3 in ARPs by CRISPR/Cas9-mediated gene editing in three independent experiments (Supplementary Figure 7). After 24h NAV3-inactivated ARPs had lost α-SMA and tubulin organization and reverted to being morphologically more similar to QuiRPs (Figure 7a). Cells were less extended and linear organization of NAV3 was no longer apparent. In liver, activated HSCs are notable for their migration to areas of tissue injury. We used live cell tracking to follow 45 ARPs over 24h. NAV3 inactivation reduced ARP migration compared to wildtype cells (Figure 7b-d). Live cell imaging over 16h showed the fluorescence intensity by reporter labelling of endogenous F-actin was also significantly reduced in NAV3-deficient ARPs (Figure 7e-f). To extend these data to human, we isolated, cultured and then abrogated NAV3 levels in human renal fibroblasts. Consistent with the requirement for active YAP1 signaling in kidney fibrosis, diminution in NAV3 increased the ratio of inactive phosphoYAP to total YAP (Figure 7g). We predicted this altered mechanosensing would alter nuclear membrane factors known to transduce alterations in cell tension and, indeed LAMIN A/C was also at least partially reduced following abrogation of NAV3. In combination, these data demonstrate NAV3 upregulation as renal pericytes transition into myofibroblasts and that NAV3 is required for the profibrotic phenotype including cytoskeletal organization and cell migration linked to the functional status of YAP1.

**Figure 6.**
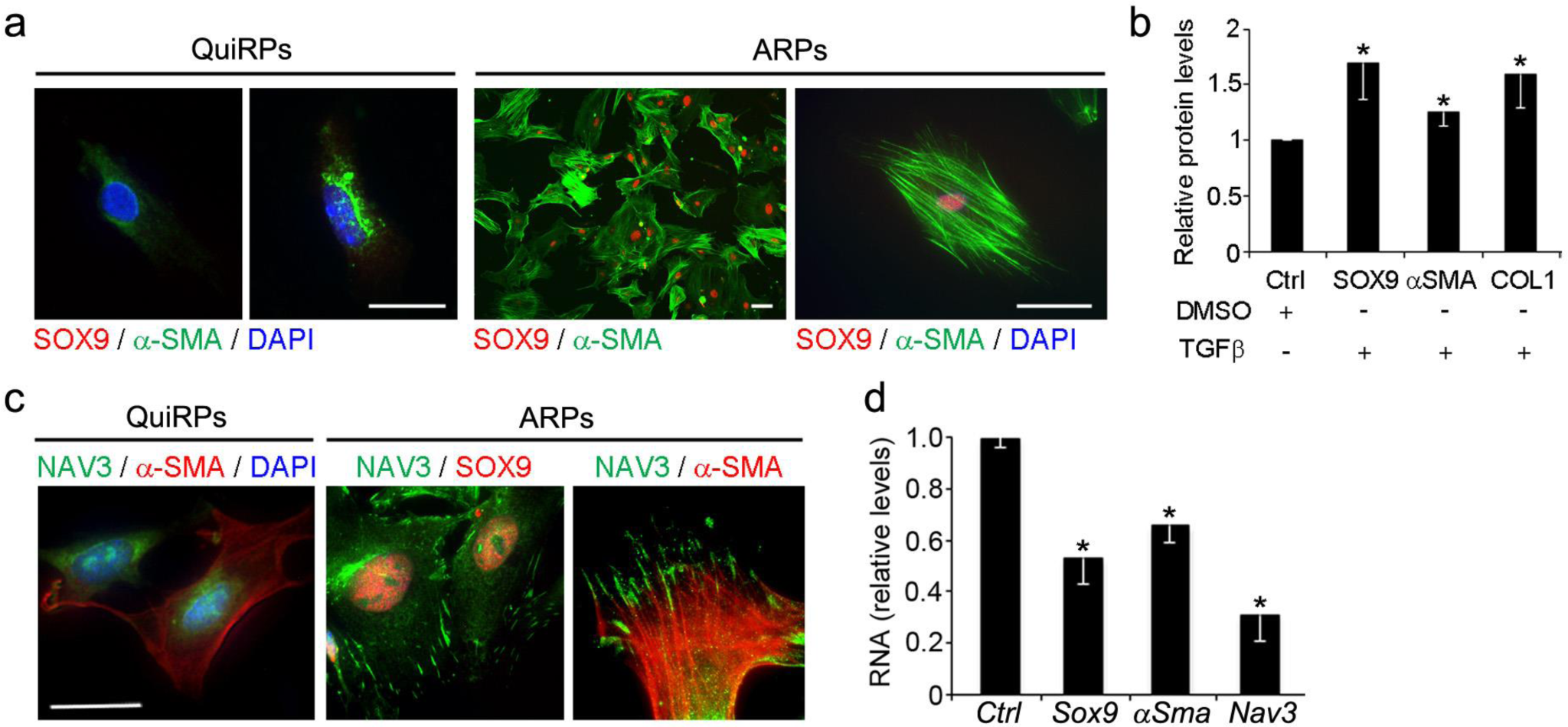
Renal pericytes transition from quiescence to activated myofibroblasts as an in vitro model of renal fibrogenesis. (**a**) Immunofluorescence of mouse renal pericytes shows shape change from rounded (quiescent, QuiRP) to myofibroblast (activated, ARP) by in vitro culture. SOX9 (red) becomes expressed and localizes to the nucleus. α-SMA (green) increases and becomes organized. (**b**) ARPs are responsive to profibrotic TGFβ signalling by increasing SOX9, α-SMA and COL1 levels. (**c**) Dual immunofluorescence for NAV3 (green) and α-SMA or SOX9 (red) counterstained by DAPI shows upregulation of NAV3 in ARPs and organisation to the ends of microtubules. (**d**) Reduction in *α-Sma* and *Nav3* relative expression by qRT-PCR following *Sox9* knockdown in ARPs. Size bars = 15 μm. Charts show means ± s.e.m, *P<0.05.

**Figure 7.**
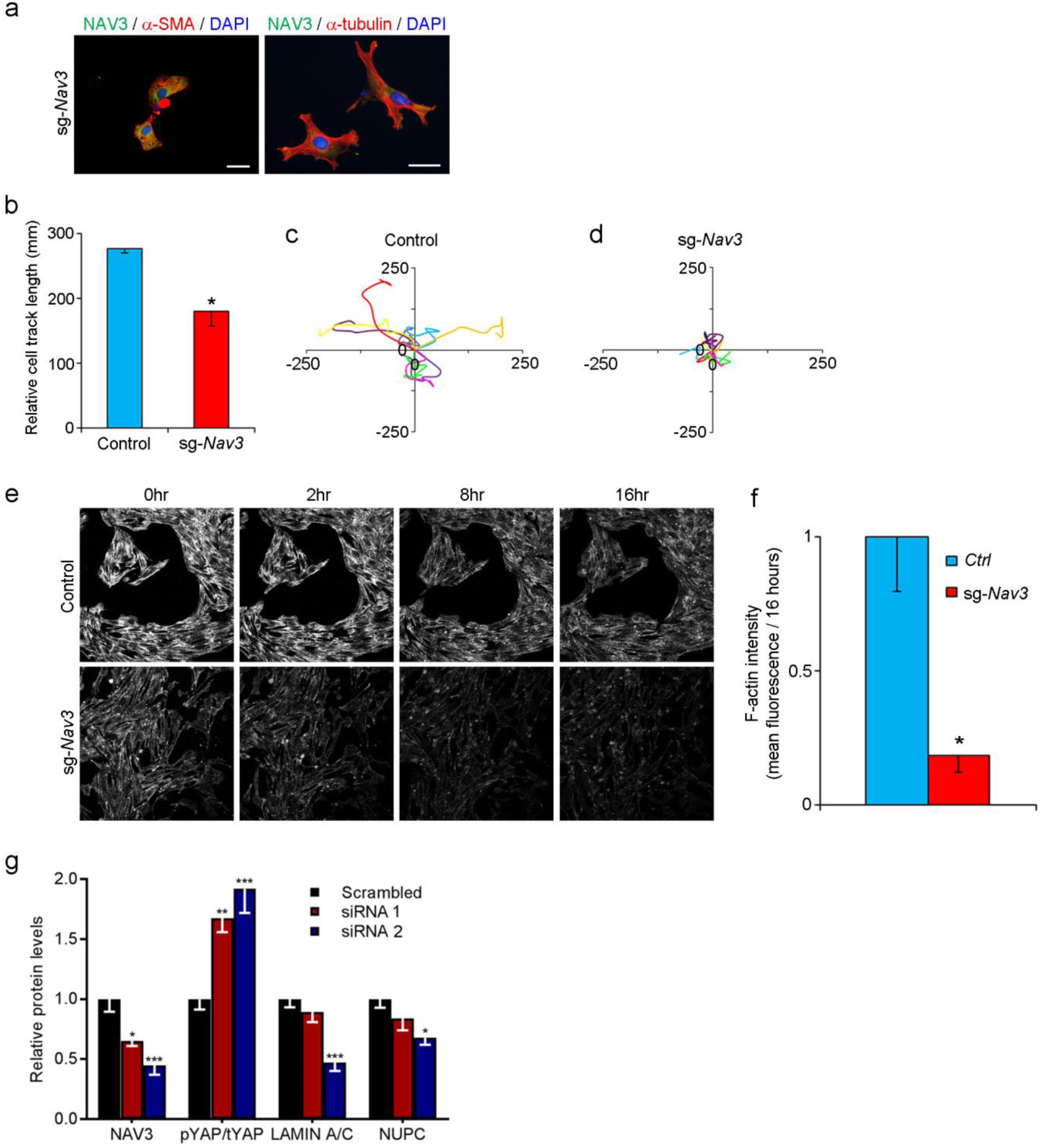
NAV3 is required for activated renal pericyte migration and maintenance of microfilaments. NAV3 was abrogated in activated renal pericytes (ARPs) by CRISPR/Cas9 using single guide sequences against *Nav3* (sg-*Nav3*). (**a**) Dual immunofluorescence for NAV3 (green), α-SMA (red) or α-tubulin (red) counterstained with DAPI shows loss of actin organization with NAV3 deficiency. (**b-d**) Migration of ARPs is attenuated by loss of NAV3. (**b**) Quantification of track length from three biological replicates (n=15 cells/experiment). (**c-d**) Example tracks in μm of different cells (coloured lines) for control ARPs (**c**) and NAV3-deficient ARPs (**d**). (**e-f**) F-actin labelling in live control or NAV3-deficient ARPs imaged over 16 h. Example images shown in (**e**) with quantification from three biological replicates (n=15 cells/experiment) in **f**. (**g**) *NAV3* knockdown by siRNAs in human renal fibroblasts increased the proportion of inactive phosphorylated YAP (pYAP) to total YAP (tYAP), and decreased protein levels for LAMIN A/C and nuclear pore complex proteins (NUPC). Two different siRNAs (siRNA2 > siRNA1) are shown compared to scrambled control. Size bar = 10 μm. Charts show means ± s.e.m, *P<0.05, **P<0.01, ***P<0.005.

## Discussion

Kidney fibrosis is a common feature that complicates many CKDs. However, our mechanistic understanding of how renal myofibroblasts arise and function is far from complete, hindering diagnosis, disease prediction and intervention. Our study helps to redress this knowledge gap by making two main discoveries, which demonstrate: the transcription factor, SOX9, is required for renal fibrosis; and, downstream of SOX9, the cytoskeletal factor, NAV3, is required by renal myofibroblasts for fibrogenesis. The findings in mouse mapped consistently to tissue from patients with chronic kidney disease.

Our observed distribution of SOX9 post-injury fits with its known role in tubule regeneration ^2, 20^. However, a series of findings collectively indicate a previously unappreciated, comprehensive role for the transcription factor in kidney fibrosis: its appearance in vitro as pericytes activate into myofibroblasts; its tissue colocalization with α-SMA and PDGFRβ in myofibroblasts; and, upon deletion, reduced expression of a profibrotic gene set, and markedly diminished interstitial collagen and lessened scarring. These new insights are consistently analogous to findings in liver fibrosis (and regeneration) where SOX9 is central to HSC activation and fibrogenesis ^7, 8, 12, 14, 15, 24-30^. The inter-organ parallels are important for at least two reasons. Firstly, the prevalence of SOX9 in tissue biopsies and serum concentration of proteins encoded from its target genes could be prognostic for progressive renal fibrosis, as we have shown in liver ^8, 12^; and secondly, equivalence with the HSC imputes the renal pericyte as an important source of myofibroblasts in interstitial kidney fibrosis. How renal myofibroblasts arise has been contentious ^31^. Our methodology here for renal pericyte-to-myofibroblast transition ought to have wide utility, including therapeutic screening, as an *in vitro* model of renal fibrogenesis.

Our genome-wide approach and differential expression analysis deciphered a broad repertoire of profibrotic SOX9 target genes. The range of genes encode ECM components including ten collagen subunits, consistent with the decreased interstitial scarring in SOX9 null kidneys post-UUO (Supplementary Figure 4). This strategy identified *Nav3*, a member of the neuron navigator gene family encoding actin-binding proteins with roles in actin remodeling ^32^. NAV3 has not been previously associated with organ fibrosis or myofibroblasts.

NAV3 has been shown to promote development and migration of the liver and pancreatic buds in zebrafish ^33^; findings which are concordant with other known SOX9 functions ^34^. We demonstrate that NAV3 is required for myofibroblast migration concordant with its stabilization of microtubules, which help guide neuronal projection ^35^. Surprisingly, in breast and hepatocellular carcinoma cells it was the opposite, where NAV3 loss-of-function increased speed of migration ^23, 36^. While superficially at odds with our findings and perhaps indicating a difference between primary and transformed cells, the more detailed study in breast cancer cells revealed the faster migration was in fact less persistent and more random ^23^. Conversely, the authors showed over-expression of NAV3 enhanced directional persistence in response to growth factors, albeit more slowly. Collectively, this would fit with our overarching concept of activating pericytes inducing SOX9 to produce, in turn, NAV3 that enables chemotactic stimuli and gradients of growth factors, such as TGF-β, to be followed to sites of renal injury. We also found that higher NAV3 levels proportionally favored more active (dephosphorylated) YAP. *SOX9* itself is a YAP target gene ^7, 8, 37^ offering an explanation for how myofibroblasts might sustain an activated phenotype in renal fibrosis via SOX9-NAV3-YAP1-SOX9 circuitry. Finally, there are no drug treatments for renal fibrosis. SOX9, as a non-liganded transcription factor, would be hard to target. In contrast, NAV3 is an ATPase. Modulating enzymatic activity is a far more tractable approach for drug development.

In summary, we have discovered a fundamental role for SOX9 and its downstream target, NAV3, in driving myofibroblast migration and function in chronic kidney disease. The data help redress the shortfall in mechanistic understanding and open the door to new therapeutic options in renal fibrosis.

## Methods

### Mouse experiments

The Sox9^fl/fl^;RosaCreER mice have been described previously ^8^. Adult (8 week) male littermates were used and genotyping was performed from DNA prepared from ear clips following our published protocol ^8^. Mice were housed and maintained, and animal experiments performed under approval from the University of Manchester Ethical Review Committee and UK Government Home Office license for animal research. 120mg/kg Tam was injected intraperitoneally (i.p.) three times in the week prior to UUO to generate Sox9 null (Sox9^fl/fl^;RosaCreER^+/−^) and Sox9 control (Sox9^fl/fl^;RosaCreER^−/−^) mice. Under isoflurane anaesthesia, the ureter of the left kidney was isolated via central laparotomy and sutured closed in 2 places to create UUO and induce ipsilateral renal fibrosis. A maintenance dose of 60mg/kg Tamoxifen was given 1 week post-UUO. Mice were sacrificed 10 days post-UUO and both kidneys removed (the right kidney, contralateral to the UUO, serving as control).

### Human tissue

Following ethical approval and informed consent, human kidney biopsy samples were acquired from the Salford Kidney Study (Regional Ethics Committee Approval 15/NW/0818; IRAS 191926; n=3 for each disease-type). Tissue collection and handling was undertaken as previously described ^8, 38, 39^.

### Histology, immunohistochemistry, immunocytochemistry and *in situ* hybridization

Picro-sirius red (PSR) was used to stain kidney sections for collagen. The extent of scarring was determined by morphometric analysis of PSR. The 3D Histech Panoramic 250 Flash II slide scanner was used to acquire images of whole kidney. At 10 x magnification 10 images were selected from each slide at random and analyzed with Adobe Photoshop. Stained pixels were selected using the Colour Range tool and expressed as a fraction of the total number of pixels.

Immunohistochemistry was undertaken as previously ^7, 8^. In brief, tissue was fixed in 4% paraformaldehyde and embedded in paraffin wax for sectioning at 5µm intervals. Heat induced epitope retrieval was performed in sodium citrate buffer, pH 6. Detection was performed using the following primary antibodies; anti-COL1 (1310-01, Southern biotech, dilution: 1:200), anti α-SMA (SMA-R-7-CE, Leica Biosystems, dilution: 1:100), anti-SOX9 (AB5535, Millipore, dilution: 1:2000), anti-NAV3 (HPA032111, Atlas Antibodies, dilution 1:2000) and anti-PDGFRβ (28E1, Cell Signalling, dilution: 1:100), followed by incubation with either Anti-Rabbit IMMPRESS AP reagent and development with ImmPACT Vector Red substrate (Vector), or incubations with species specific biotinylated antibodies (Vector, 1:100), then streptavidin-HRP antibody (Vector, 1:200) before development with diaminobenzidine (Sigma). Counterstaining was performed using toluidine blue. Picrosirius red staining and quantification of staining was performed as previously described ^7, 8^.

For immunocytochemistry, cells were cultured on glass chamber slides (Lab-Tek), fixed in 4% paraformaldehyde for 10 minutes and incubated with primary antibodies. For all immunofluorescence secondary antibodies were 488 or 594 Alexa Fluors (Molecular Probes, Invitrogen) raised against the appropriate species (dilution: 1:1000).

*In situ* hybridization of target transcripts in paraffin embedded sections was performed using a multiplex fluorescent RNAScope assay (ACD) on a BOND RX automated staining system (Leica Biosystems) according to manufacturer’s instructions using probes against *Nav3* (Cat No. 539128), *Acta2* (Cat No. 319538) and *Sox9* (Cat No. 401058). Sections were mounted using Vectashield with DAPI (Vector) prior to imaging with a 3D Histech Pannoramic 250 Flash Slide Scanner (Zeiss) with a 40x objective lens. *Nav3* mRNA quantification was performed in 20 fields of view for each of the 4 groups (Sox9 null and Sox9 control mice / UUO kidney and contralateral unaffected kidney; n=3 per group, 12 samples in total).

All quantification was carried out blind to the investigators following randomization of slides and acquired images.

### Pericyte preparation

A modified version of the protocol in Schrimpf *et al* (2012) was used ^40^. Briefly, the renal capsule was removed from both kidneys before mincing with a razor blade and digestion at 37°C for 30 min in HBSS+ (ThermoFisher) containing 0.8U/ml Liberase TM (Roche) and 1mg/ml DNAse (Sigma). Digestion was stopped with the addition of 10% FBS and the suspension filtered until all debris was removed using sequential 100µm (x1), 70µm (x1) and 40µm (x2) sterile nylon filters. After two centrifugation (300g) and wash cycles in 10% FBS in DMEM, the sample was centrifuged a final time, the supernatant discarded, and the cell pellet resuspended in de-gassed MACS solution (Miltenyl Biotech). Cells were incubated with anti-rabbit Biotin-PDGFRβ (Miltenyi, Dilution 1:11) for 15 mins on ice before two centrifugation and washing cycles in cold MACS buffer. Cells were then incubated with goat anti-rabbit IgG microbeads (Miltenyi biotech, Dilution 1:20) for a further 15 mins on ice before three further cycles of centrifugation and washing in cold MACS buffer and resuspension in MACS buffer. Perictyes were captured on LS columns (Miltenyi Biotech) inserted into a quadraMACS magnet (Miltenyi Biotech). After removal from the magnet, cells were eluted from the column for culture in DMEM/F12 (ThermoFisher) enriched with 1% L-Glutamine, 10% FBS, 1% Pen/Strep and 1% Insulin-Transferrin-Selenium (ITS).

### qRT-PCR

Tissue was stored in RLT buffer at −80°C until use. RNA was extracted using an RNeasy kit (Qiagen), followed by DNAse I treatment (Sigma) and conversion into cDNA using a High Capacity RNA-to-cDNA kit (Life Technologies). Assays were performed using primers (Supplementary table 2) and a SYBR green master mix (Primerdesign) on a StepOnePlus system (Applied Biosciences). Ct values were standardized to two ‘housekeeping’ genes, *GusB* and *ActinB*, and relative quantification calculated using ΔΔCt methodology.

### RNA sequencing

RNA was prepared ^7^ and RNAseq undertaken as previously described ^41^. RNA quality checks, creation of a cDNA library and DNA fragmentation was performed by the University of Manchester Genomics Core Facility. RNAseq was undertaken in biological replicate using the Illumina HiSeq 4000 to generate 280-300 million reads across ipsilateral UUO and contralateral unaffected kidneys from Sox9 null and Sox9 control mice (8 samples total). Post-sequencing QC analysis was undertaken as previously ^41^. In brief, paired-end sequences were tested by FastQC (http://www.bioinformatics.babraham.ac.uk/projects/fastqc/), sequence adapters removed and reads quality trimmed using Trimmomatic_0.36 ^42^. Reads were mapped against the reference mouse genome (mm10/GRCm38) and counts per gene were calculated according to annotation from GENCODE M21 (http://www.gencodegenes.org/) using STAR_2.5.3 ^43^. Normalization, principal components analysis, and differential expression was undertaken with DESeq2_1.24.0 ^44^. Cluster analysis was performed using a κ-means clustering algorithm to place gene expression into groups. Pathways and networks associated with the differential gene expression were analyzed using Ingenuity Pathway Analysis (Ingenuity® Systems, www.ingenuity.com).

### Western blot

Quantification of protein by Western blotting was carried out using standard techniques and the following primary and secondary antibodies: anti-α-SMA (SMA-R-7-CE, Leica Biosystems, dilution: 1:100), anti-COL1 (1310-01, Southern Biotech, dilution:1:1000), anti-SOX9 (AB5535, Millipore, dilution: 1:5000), anti-YAP (Sc-271134, Santa Cruz, dilution: 1:1000), anti-NAV3 (ab69868, Abcam, dilution 1:100) anti-Pdgfrβ (28E1, Cell Signalling, dilution 1:1000), anti-pYAP1 (4911, Cell Signalling, dilution 1:1000), anti-YAP1 (4912, Cell Signalling, dilution 1:1000), anti-NUPC (24609, Abcam, dilution 1:1500), anti-Lamin A/C (2032, Cell Signalling, dilution 1:1000), anti-β-actin HRP conjugate (A3854, Sigma, dilution: 1:50000) and species-specific HRP conjugated secondary antibodies (GE Healthcare). ECL Advanced Western blot detection reagent (GE Healthcare) was applied to the membrane and signal intensity measured on a Gel Doc Imager (BioRad). Data were normalized to GAPDH expression or total protein determined via UV treatment of TGX Stain-Free gels (Biorad) followed by detection on the transfer membrane.

### CRISPR/Cas9 editing of *Nav3* in mouse renal pericytes

Target single guide (sg) sequences in the *Nav3* gene were identified in exon 1 (5’-gacacactgatgccaagatt-3’) and exon 2 (5’-gcagcaggacattgcggacg-3’) using E-CRISP ^45^. Two *Nav3*-targeting plasmids were generated by the insertion of annealed, phosphorylated oligonucleotides containing each target sequence into *Bbs*I-digested pSpCas9(BB)-2A-Puro (PX459) V2.0 (Addgene plasmid # 62988) ^46^. Mouse renal pericytes were co-transfected with both plasmids with Lipofectamine 3000, or empty PX459 as control. Transiently transfected cells were selected as a total population with Puromycin-enriched media (0.5 μg/ml) for 48 h. To confirm effective mutagenesis, a high percentage PAGE assay was used on PCR products amplified from the target site of selected cells to detect heteroduplex formation (adaptation of ^47^). Loss of *Nav3* was confirmed via qRT-PCR and Western blot. Accordingly, *Nav3* CRISPR/Cas9 pericytes refers to a transiently transfected population of cells treated with short-term puromycin.

### siRNA knockdown of *NAV3* in HEK293 cells

Gene silencing in HEK293 cells was carried out via transient transfection of 10 nmol siRNA (*NAV3*_siRNA1: SI04272646, *NAV3*_siRNA2: SI04366215) or scrambled control (Qiagen) using Lipofectamine 3000 (ThermoFisher). Cells were harvested 48 h post-transfection in RIPA buffer for Western blot.

### Luciferase assay

Conserved SOX9 binding motifs surrounding the human *NAV3* locus were identified using Mulan alignment (https://mulan.dcode.org/) and multiTF (https://multitf.dcode.org/) ^48^. Approximately 100bp of the surrounding sequence was amplified by PCR from LX2 cells and cloned into the *Xho*I site of the pGL3+promoter vector (Promega) with corresponding empty vector as control. Mutated amplicons lacking the conserved ACAAT SOX9-binding motifs were also cloned into the same vector (coordinates and sequences in Supplementary Figure 6 and Supplementary Table 3). LX2 cells were transfected using Lipofectamine 3000 with the reporter plasmids, ‘*Renilla*’ pRL-TK (Promega), and pcDNA3.1Zeo+ empty vector or pcDNA3.1Zeo+ containing a full length human SOX9 coding sequence. Vector inserts were confirmed by sequencing. 48 h post-transfection, luciferase activity was measured in cell lysates using the Dual-Luciferase reporter assay system (Promega) normalized to *Renilla* luciferase activity.

### Cell migration

*Nav3* CRISPR/Cas9 and control renal pericytes were cultured in 24-well plates at a density of 5,000 cells per well. Single-cell tracks were acquired using live-cell imaging at 10 min intervals over 24 h (AS MDW live cell imaging system, Leica). Multiple position imaging was gathered using Image Pro 6.3 (Media Cybernetics Ltd) and a Coolsnap HQ (Photometrics) camera with a Z optical spacing of 0.2 mm. Cell movement over 24 h was analyzed using ImageJ software with the MTrackJ plug-in. Total length of the cell track (in mm) was generated for multiple cells, averaged and normalized to control. Where individual migration tracks are shown, co-ordinates at 10 min intervals were plotted relative to the cell’s origin.

### Live F-actin staining

10,000 *Nav3* CRISPR/Cas9 or control renal pericytes were seeded into the 6 central wells of a 24-well plate. 1µl/ml SiR-actin (CY-SC001, Spirochrome) was added per well and live cell imaging commenced following exchange after 2h into fresh media containing 1µl/ml Verapamil to enhance SiR-actin staining. SiR-actin intensity was measured at fifteen coordinates for control and *Nav3* CRISPR/Cas9 cells using a CSU-X1 spinning disc confocal (Yokagowa) on a Zeiss Axio-Observer Z1 microscope with a 20x objective. Slidebook software (3I) was used to capture images every 10 min over 16 h. LASX software (Leica) calculated net intensity from each co-ordinate. Statistical significance for mean intensities per well was determined between the two groups (*Nav3* CRISPR/Cas9 and control) by two-tailed unpaired T-test. Cells were incubated at 37 °C and 5% CO_2_ for the duration of the experiment.

### Statistics

All statistical analyses were performed using GraphPad Prism 7 (GraphPad). Two-tailed T-tests were used for single comparisons. For multiple comparisons, 1-way or 2-way ANOVA was performed with Dunnett’s post-hoc multiple comparisons test to identify the significance of individual comparisons.

## Author Contributions

KPH and NAH conceived and designed experiments. SR and EJ contributed to experimental planning and design. NCH and PAK provided reagents and contributed to experimental design. SR, EJ, KM, KS, AFM, KS, VA and JP performed experiments. SR, EJ and KPH analyzed the data. LZ carried out bioinformatics and statistical analysis. KPH guided experiments, analyzed data and wrote the manuscript.

## Acknowledgements

This work was supported by the Medical Research Council (MRC; KPH, MR/J003352/1 & MR/P023541/1; NAH, MR/000638/1 & MR/S036121/1; and KM and VSA are MRC Clinical Training Fellows). NAH was a Wellcome senior fellow. SR is a KRUK Clinical Training Fellow and received funding support for this work by Kidneys for Life. The Genomics Core Facility and the Bioimaging Facility at the University of Manchester are acknowledged for their technical help and support from Wellcome (105610).

