## Supplementary data for "SOX9 is required for kidney fibrosis and regulates NAV3 to control renal myofibroblast function in mice and humans"

***This document includes Supplementary Figs. 1-7 and Supplementary Tables 2-3. Supplementary Table 1 is available as a separate Excel file.**

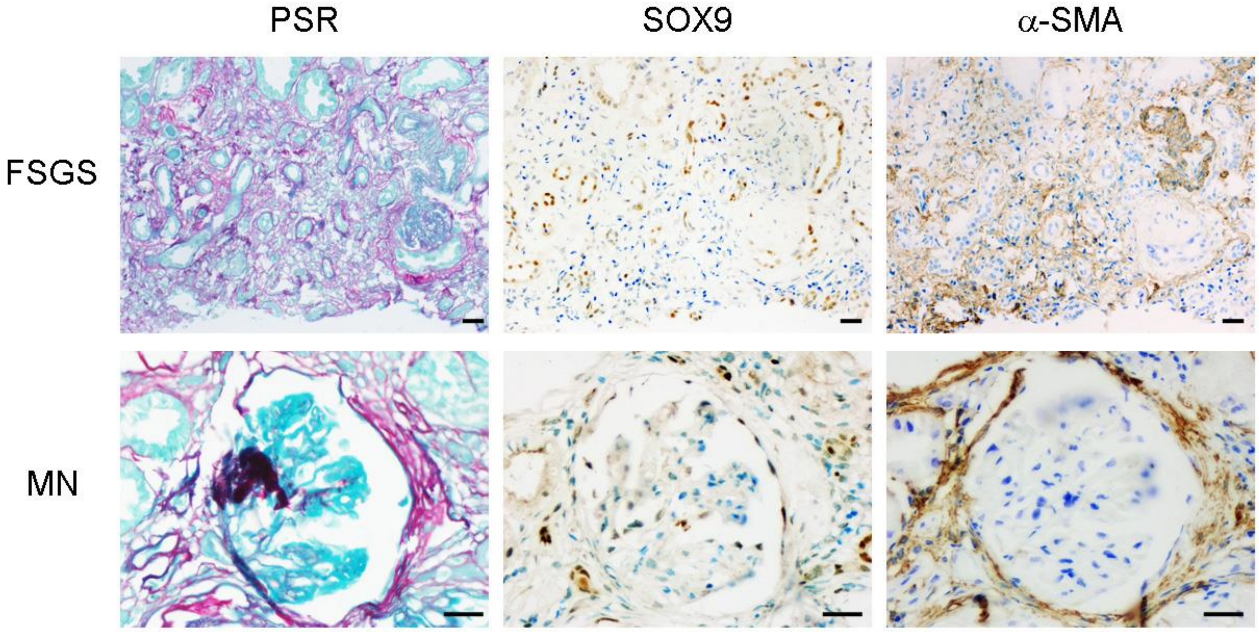


**Supplementary Figure 1. Localization of SOX9 in different forms of chronic kidney disease in patients.** Serial kidney sections from focal segmental glomerulosclerosis (FSGS; top panel) and membranous nephropathy (MN; bottom panel) showing collagen deposition by picrosirius red (PSR; red) staining, and immunohistochemistry for SOX9 (brown; middle) and α-SMA (brown; right). Nuclear SOX9 localizes to tubules and interstitial areas with α-SMA-positive myofibroblasts. Size bar = 25µm.

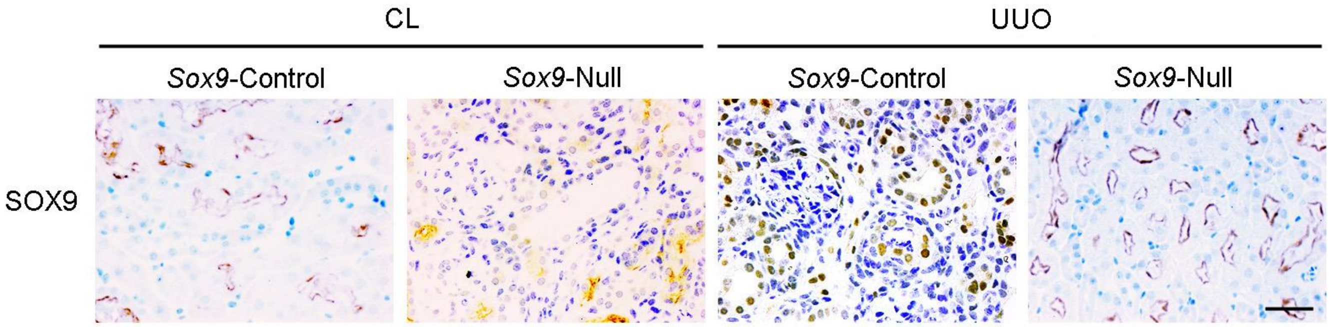


**Supplementary Figure 2. SOX9 loss following recombination in Sox9 null mice (refers to Fig. 3).** Representative images shown for *Sox9* null and *Sox9* control mice in the ipsilateral kidney following UUO or unaffected contralateral (CL) kidney (n=4 for all groups). Immunohistochemistry for SOX9 (brown) is counterstained with toluidine blue. Note the non-specific staining in the brush border of kidney tubules in each panel. However, no nuclear SOX9 is detected in the CL kidneys of *Sox9* control mice (i.e. although the *Sox9* gene is intact nuclear SOX9 is not present in the absence of renal injury). Nuclear SOX9 is detected following UUO in the ipsilateral kidney of *Sox9* control mice but not *Sox9* null mice (indicating the efficacy of *Sox9* deletion; far right panel). Size bar = 100 µm.

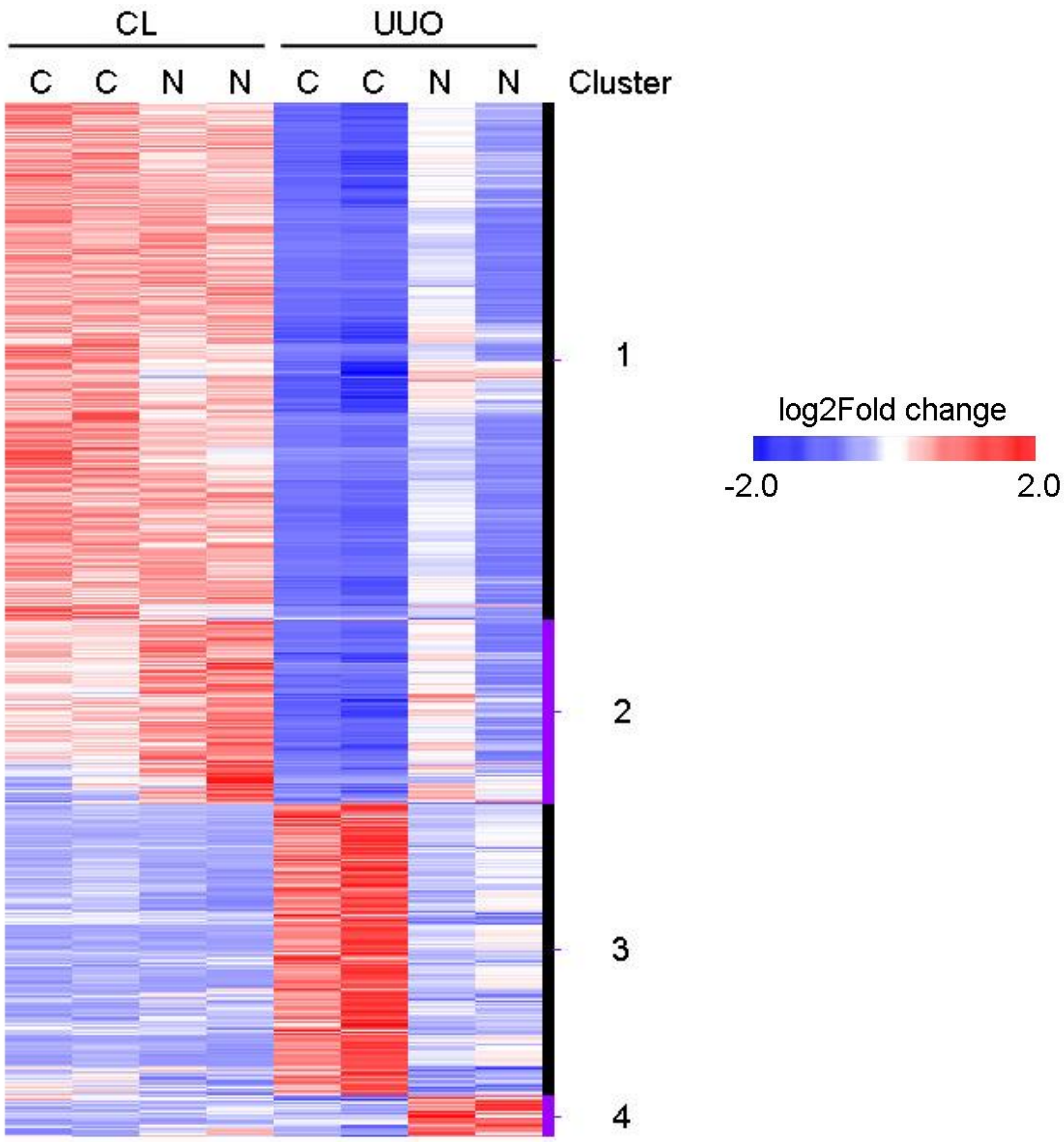


**Supplementary Figure 3. Heatmap of hierarchical clustering of RNAseq data.** Data are displayed by heatmap for those genes with log2-fold differences in expression >2.0. Both biological replicates are shown from which the mean data was obtained as shown in Fig. 4a. Genes are filtered for statistical significance ($P<0.05$). *Sox9* control mice (C); *Sox9* null mice (N); contralateral unaffected kidney (CL); and ipsilateral kidney injured following UUO. Four clusters were identified based on upregulated (red) and downregulated (blue) gene expression.

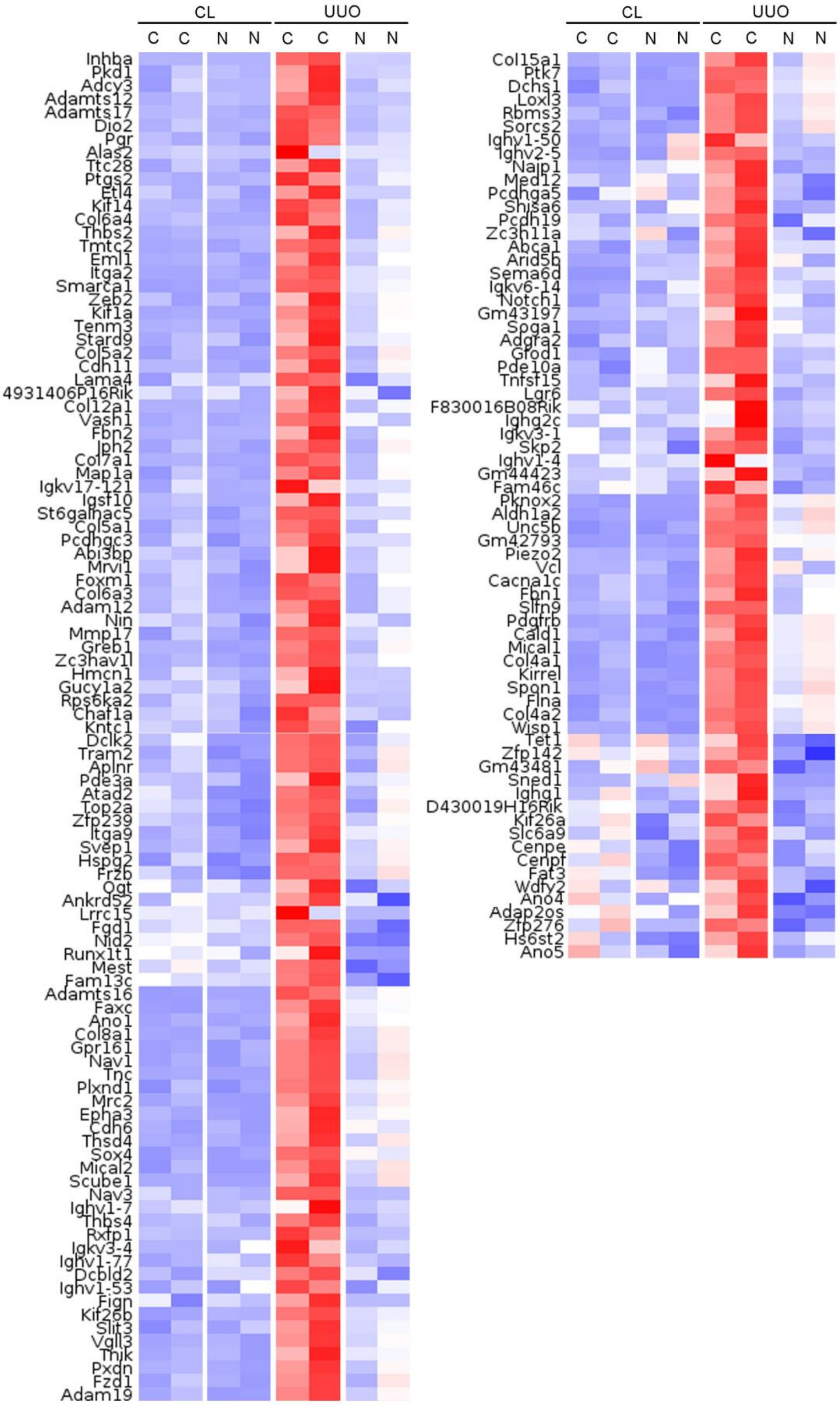


**Supplementary Figure 4. Heatmap of hierarchical clustering for Cluster 3 showing full gene list.** The layout and abbreviations are identical to Supplementary Figure 3. The data relate to Fig. 4a of the main manuscript. Color indicates upregulated (red) and downregulated (blue) gene expression.

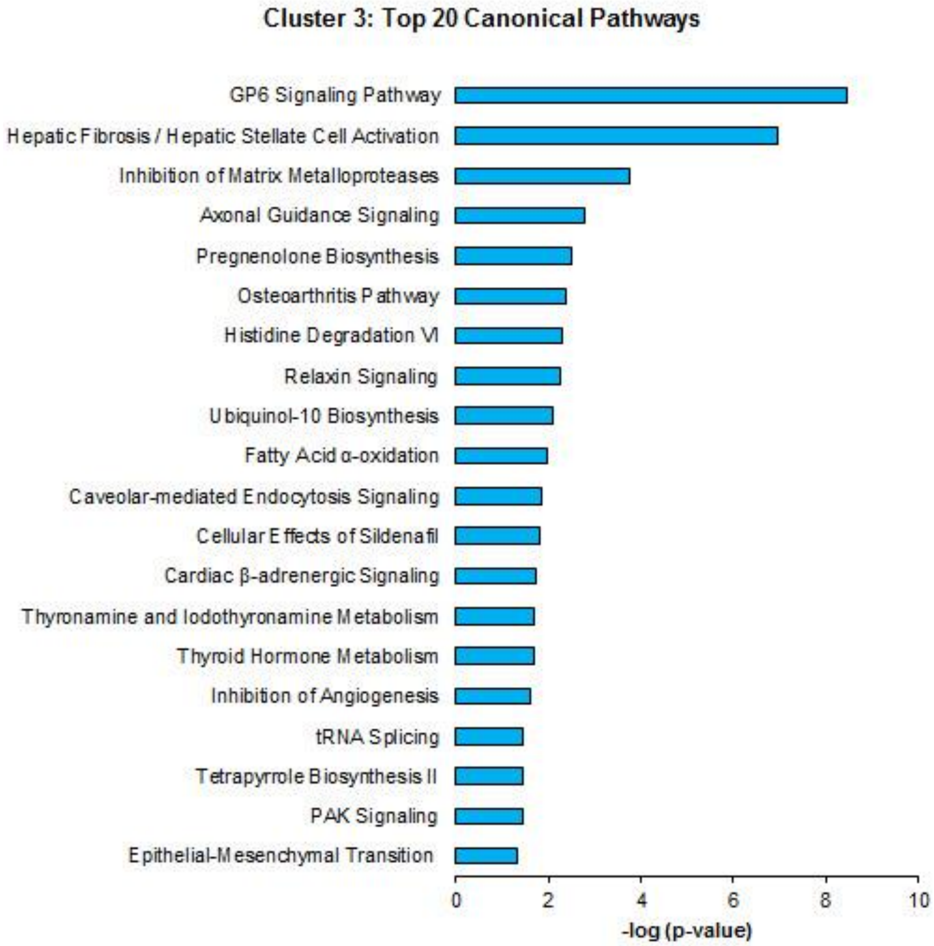


**Supplementary Figure 5. Top 20 canonical pathways for genes enriched in Cluster 3 by Ingenuity Pathway Analysis 3.** See Fig. 4a and Supplementary figure 4. Pathways were ranked by the negative log of P-values calculated by Fisher's exact test for gene enrichment.

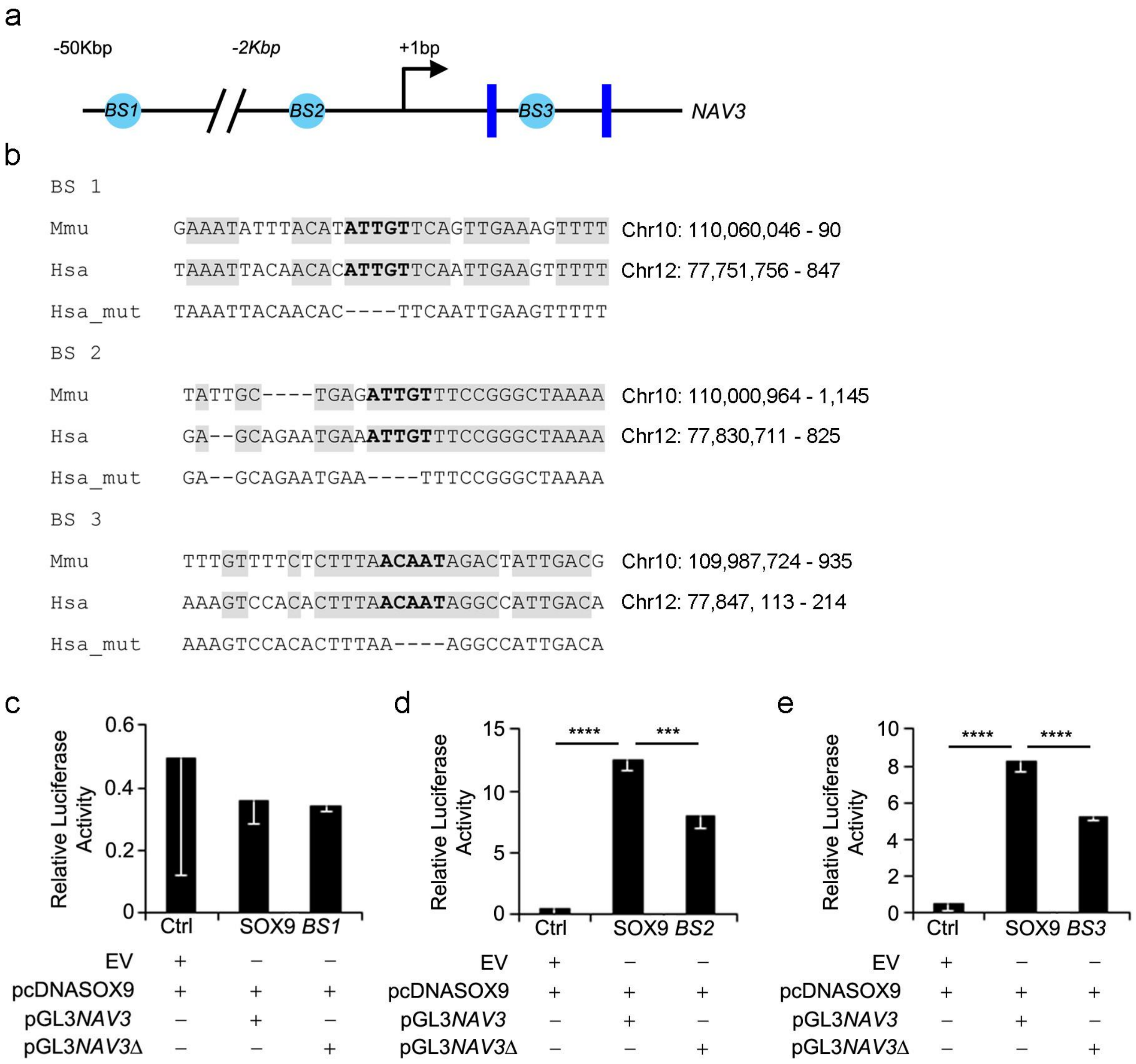


**Supplementary Figure 6.** (**a**) Schematic showing conserved SOX9 binding sites in *NAV3*. (**b**) Sequence showing conserved SOX9 binding sites in bold, conservation across mouse (Mmu) and human (Hsa) in grey, and the mutated sites in the human sequence (Hsa_mut) for binding sites 1-3 (BS1-BS3). (**c-e**) Luciferase activity (in relative light units) following transient transfection with indicated constructs for BS1 (**c**), BS2 (**d**) and BS3 (**e**). In each instance, pGL3*NAV3* and pGL3*NAV3*Δ represent the wildtype and mutated SOX9 binding site respectively in the human sequence. Data in bar charts show means ± s.e.m. ***P<0.005, ****P<0.001.

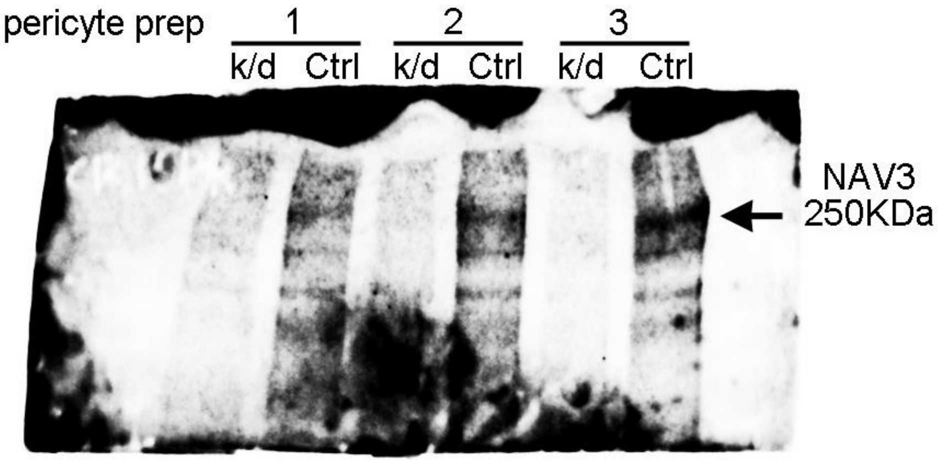


**Supplementary Figure 7. Immunoblotting and verification of NAV3 abrogation following CRISPR/Cas9 for Figure 7 of the main manuscript.** Three individual pericyte preparations are shown. NAV3 protein indicated for knockdown (k/d) cell lysate (corresponding to sg-*Nav3* from Fig. 7 main text) and control lysate (Ctrl).

| **Target gene** | **Forward Primer (5’ to 3’)** | **Reverse Primer (5’ to 3’)** |
|---|---|---|
| *Ppia* | *gagctgtttgcagacaaagttc* | *ccctggcacatgaatcctgg* |
| *Sox9* | *ggccgaagaggccacggaac* | *gattgcccagagtgctcgccc* |
| *Acta2* | *gtcccagacatcagggagtaa* | *tcggatacttcagcgtcagga* |
| *Col1a1* | *gctcctcttaggggccact* | *ccacgtctcaccattgggg* |
| *Pdgfrb* | *tccaggagtgataccagcttt* | *caggagccataacacggaca* |
| *Nav3* | *tccgataccgaatcttggcat* | *ggcacgaaagcattgagcg* |
| *GusB* | *gcagttgtgtgggtgaatgg* | *gggtcagtgtgttgttgatgg* |
| *Actb* | *gctgtattcccctccatcgtg* | *cacggttggccttagggttcag* |

**Supplementary Table 2.** Primer sequences used for RTqPCR.

| Insert | Sequence |
|---|---|
| Wildtype (WT) binding site (BS) 1 | GTGAGCATATACATTTTGTATTCTTAAATTACACAATACAATTTACACATTGTTCAATTGA AGTTTTTTTCTTTCATAACCGAGCTTATACG |
| Mutated BS 1 | GTGAGCATATACATTTTGTATTCTTAAATTACACAATACAATTTACAC---- TTCAATTGAAGTTTTTTTCTTTCATAACCGAGCTTATACG |
| WT BS 2 | GCTGTCAGTAGTGAAAAATAGCTGGAAATCAGACAAACAACTTTATTGCTGAGATTGTTTC CGGGCTAAAAGTTCTTCCAACAGCTGTTTGTTTTGGCCATTAACATGTCCATTC |
| Mutated BS 2 | GCTGTCAGTAGTGAAAAATAGCTGGAAATCAGACAAACAACTTTATTGCTGAG---- TTCCGGGCTAAAAGTTCTTCCAACAGCTGTTTGTTTTGGCCATTAACATGTCCATTC |
| WT BS 3 | CTCCTCTGTAAGACAGACAGGATGTGGTCTAGGAGAAAGTCCACACTTTAACAATAGGCCA TTGACAGAATGTGTGGGGAGTACAAAGAGGAAGAGAGCATC |
| Mutated BS 3 | CTCCTCTGTAAGACAGACAGGATGTGGTCTAGGAGAAAGTCCACACTTTAA---- AGGCCATTGACAGAATGTGTGGGGAGTACAAAGAGGAAGAGAGCATC |

**Supplementary Table 3. Full insert sequences for luciferase assay plasmids.**
